# Effects of transcranial focused ultrasound on human primary motor cortex using 7T fMRI

**DOI:** 10.1101/277608

**Authors:** Leo Ai, Priya Bansal, Jerel K. Mueller, Wynn Legon

## Abstract

**Background:** Transcranial focused ultrasound (tFUS) is a new non-invasive neuromodulation technique that uses mechanical energy to modulate neuronal excitability with high spatial precision. tFUS has been shown to be capable of modulating EEG brain activity in humans that is spatially restricted, and here, we use 7T MRI to extend these findings. We test the effect of tFUS on 7T BOLD fMRI signals from individual finger representations in the human primary motor cortex (M1) and connected cortical motor regions. Participants (N = 5) performed a cued finger tapping task in a 7T MRI scanner with their thumb, index, and middle fingers to produce a BOLD signal for individual M1 finger representations during either tFUS or sham neuromodulation to the thumb representation.

**Results:** Results demonstrated a statistically significant increase in activation volume of the M1 thumb representation for the tFUS condition as compared to sham. No differences in percent BOLD changes were found. This effect was spatially confined as the index and middle finger M1 finger representations did not show similar significant changes in either percent change or activation volume. No effects were seen during tFUS to M1 in the supplementary motor area (SMA) or the dorsal premotor cortex (PMd).

**Conclusions:** Single element tFUS can be paired with high field MRI that does not induce significant artifact. tFUS increases activation volumes of the targeted finger representation that is spatially restricted within M1 but does not extend to functionally connected motor regions.

## Introduction

Transcranial focused ultrasound (tFUS) is a noninvasive, low energy technique that uses mechanical energy for neuromodulation at high spatial resolutions [1]. tFUS has been shown to be capable of modulating neural activity in mice [2-4], rabbit [5], swine [6], and monkeys [7]. tFUS has also been shown to be a safe and effective method to modulate human cortical activity [1, 8-11]. In Legon et al. (2014), we demonstrated the spatial selectivity of tFUS neuromodulation though the spatial resolution of EEG is not ideal for this. The pairing of tFUS with functional MRI is advantageous as it provides complimentary high spatial resolution with whole brain coverage. Previous reports have shown ultrasound to elicit a blood oxygen level dependent (BOLD) response. In craniotomized rabbits, Yoo et al. (2011) showed focused ultrasound directed at the somatomotor area to result in a well-defined BOLD response commensurate with the focus of sonication. In a recent study in humans, Lee et al. (2016) delivered focused ultrasound to the primary visual cortex and showed BOLD activity around the sonication focus in visual cortices but also for ultrasound to activate spatially distinct functionally connected regions of the visual system. We have also previously tested the ability of tFUS to produce a reliable BOLD signal in humans at 3T and report variable effects (Ai et al. 2016). Here, we extend these findings and pair tFUS with high field 7T fMRI in humans to improve signal to noise ratios and the ability to discriminate small spatially restricted changes in activity from tFUS. Specifically, we apply tFUS to the human primary motor cortex (M1) and test the effect of tFUS on specific finger BOLD signals as well as on functionally connected regions including the supplementary motor area (SMA) and dorsal premotor cortex (PMd).

## Methods

### Participants

Five participants (ages 20-25 (Mean 22.8 ± 2.2 years); 3 male, 2 females; 4 right handed, 1 left handed) were included in the study. This study was approved by University of Minnesota’s Institutional Review Board and all Participants gave written informed consent to participate. Participants were physically and neurologically healthy and had no history of neurological disorders. Participants were also screened for medications contraindicated for other forms of non-invasive neuromodulation [12].

### Experimental procedures

The study consisted of two MRI scanning sessions on separate days. The first session included a T1 anatomical scan and a functional scan with the finger tapping task (see below) to identify M1 thumb, index and middle finger representations. The thumb representation was then used as the target for the application of tFUS for the second session. In the second session, participants performed the same finger tapping task during either tFUS or sham neuromodulation. The order of tFUS and sham conditions was counterbalanced across participants.

### Finger tapping task

Participants performed a visually cued finger tapping task using either the thumb, index, and middle fingers with their self-reported dominant hand. Participants lay supine in the MRI with their dominant arm supported with foam to ensure a comfortable position to tap their fingers on their thigh while limiting proximal arm and shoulder movement. Visual cues indicating the timing for tapping were presented using Cogent (www.vislab.ucl.ac.uk/cogent.php) for Matlab (MathWorks, Natick, MA, USA) and delivered using a projector to a screen that participants could see while inside of the bore of the MRI machine. The visual cues displayed the text (’thumb‘, ‘index‘, or ‘middle‘) with white block letter on a black background in the center of the screen with a large font, indicating the finger to be tapped paced at 1Hz. This task used a block design with a single finger to be tapped for the duration of a block at the 1 Hz pace. Each finger was tapped for three blocks for a total of nine 30 second blocks, with 30 second rest blocks separating each finger tapping block (Figure 1A). The ordering for the finger to be tapped per block was pseudo-randomly generated for each MRI scan where no finger would be tapped for three contiguous blocks.

**Figure 1.**
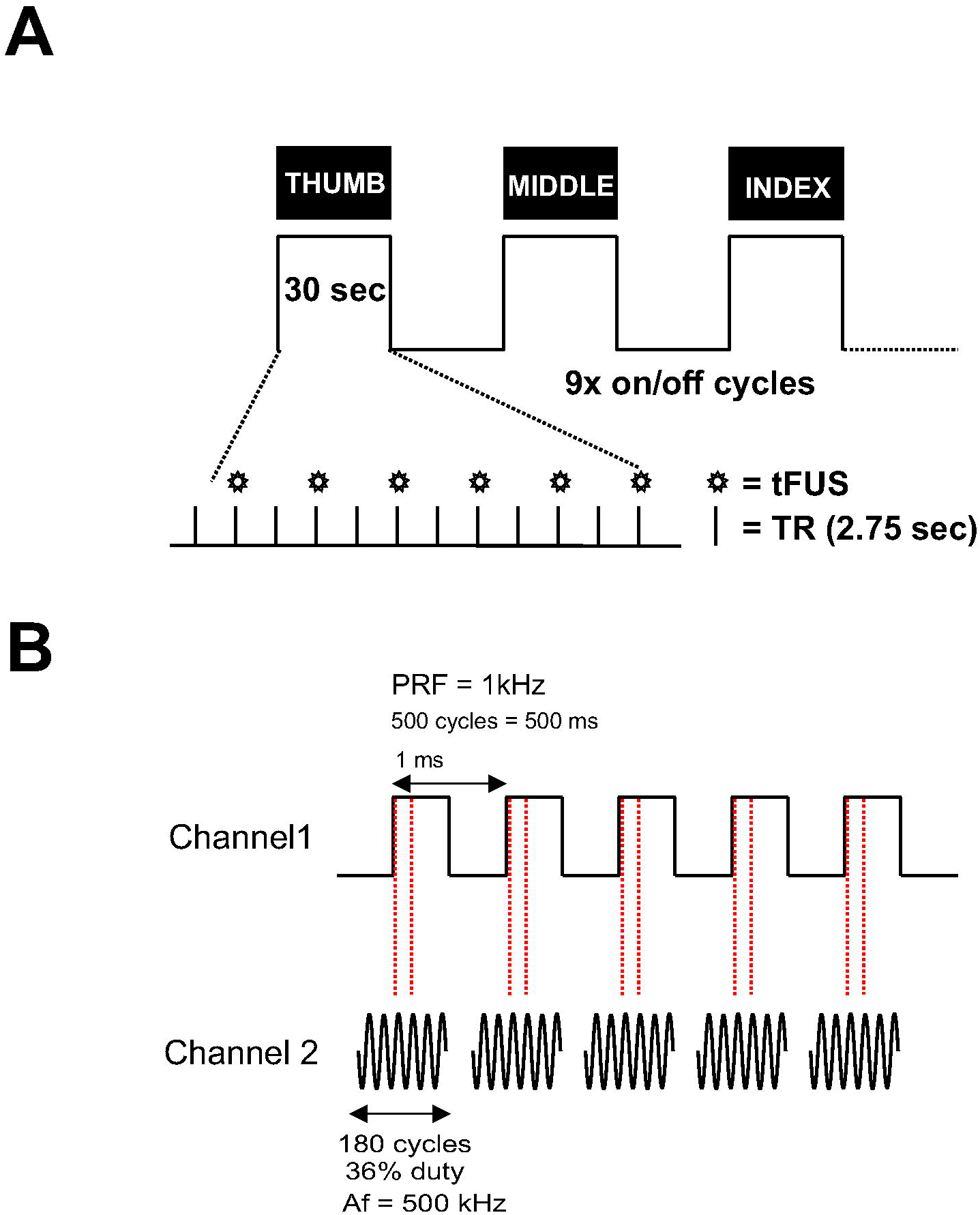
**A**. Schematic of the fMRI experimental protocol. Finger movement (thumb, middle, index) was visually cued at 1Hz across the on blocks. A total of nine 30 second on blocks were collected (3 for each finger) interspersed with 30 second rest blocks. Within each on block transcranial focused ultrasound (tFUS) was delivered every two TRs. **B.** Schematic of the ultrasound pulsing strategy. PRF = pulse repetition frequency; Af = acoustic frequency.

Prior to scanning, participants practiced the finger tapping task to familiarize themselves with the task demands. To standardize movement range, participants were instructed to follow the visual prompts by extending and flexing the cued finger at the proximal phalanx while limiting movement of other fingers. Participants performed this practice session with feedback from the study staff to ensure the task would be performed properly while inside the scanner. Ultrasonic waveforms were delivered every two repetition times (TR, 2750ms) for a total of 6 stimulations per 30 second block (54 total stimulations per scan). The tFUS condition involved acoustically coupling the active face of the ultrasound transducer to the scalp at the pre-determined neuronavigation (see below) site. To achieve acoustic coupling to the head, the volunteer’s hair was parted to expose the scalp and ultrasound gel was used to keep the hair out of the way and ensure proper coupling with the tFUS transducer. The transducer was also prepped with ultrasound gel on the surface that met the head, and was then placed on the exposed scalp and held in place using a secure head band [1, 9, 10]. The sham condition involved turning off the transducer so that it would not deliver stimulation. Participants reported no auditory or tactile sensation from either the tFUS or sham condition.

### tFUS waveform and delivery

The ultrasound transducer was a custom made [13] 30 mm diameter 7T MRI compatible single element focused 500 kHz with a focal length of 30 mm. The waveform used was the same as previously described [1]. This waveform was generated using a two-channel 2-MHz function generator (BK Precision Instruments, CA, USA). Channel 1 was set to deliver tFUS at a pulse repetition frequency (PRF) at 1 kHz and channel 2 was set to drive the transducer at 500 kHz in burst mode while using channel 1 as the trigger for channel 2. Channel 2 was set to deliver 180 cycles per pulse, and channel 1 was set to deliver 500 pulses, resulting in a 500ms duration (Figure 1B). Channel 2 output was sent to a 100W linear amplifier (2100L Electronics & Innovation Ltd, NY, USA), with the output of the amplifier sent to the custom made tFUS transducer while using a Mini-Circuits (New York City, NY) 50-ohm low pass filter (1.9MHz cutoff frequency) and matching network.

### Quantitative acoustic field mapping

The acoustic intensity profile of the waveform was measured in an acoustic test tank filled with deionized, degassed, and filtered water (Precision Acoustics Ltd., Dorchester, Dorset, UK). A calibrated hydrophone (HNR-0500, Onda Corp., Sunnyvale, CA, USA) mounted on a motorized stage was used to measure the acoustic intensity profile from the ultrasound transducer in the acoustic test tank at a 0.5 mm spatial resolution. Intensity parameters were derived from measured values of pressure using the approximation of plane progressive acoustic radiation waves. The ultrasound transducer was positioned in the tank using opto-mechanical components (Edmund Optics Inc., Barrington, NJ and Thorlabs Inc., Newton, NJ). Acoustic field scans were performed in the free water of the tank. Measurements in the acoustic tank revealed an spatial peak pulse average intensity (I_sppa_) of 16.95 W/cm^2^ and a mechanical index (MI) of 0.97 from the ultrasonic neuromodulation waveform in water. The -3dB pressure field was 3.83 mm in the X axis, 3.98 mm in the axis and 33.6 mm in the Z axis (Figure 2). We have previously modelled the acoustic field through human skulls overlying the motor cortex demonstrating the skull to reduce peak pressure produced by the transducer in free water by a factor of 6 to 7, and it can be expected for the targeted region of the brain to experience pressure to be reduced as such [14]. In addition, the brain tissue and skull do not alter the beam path significantly [14, 15] or result in appreciable heating of the skin or skull bone [16].

**Figure 2.**
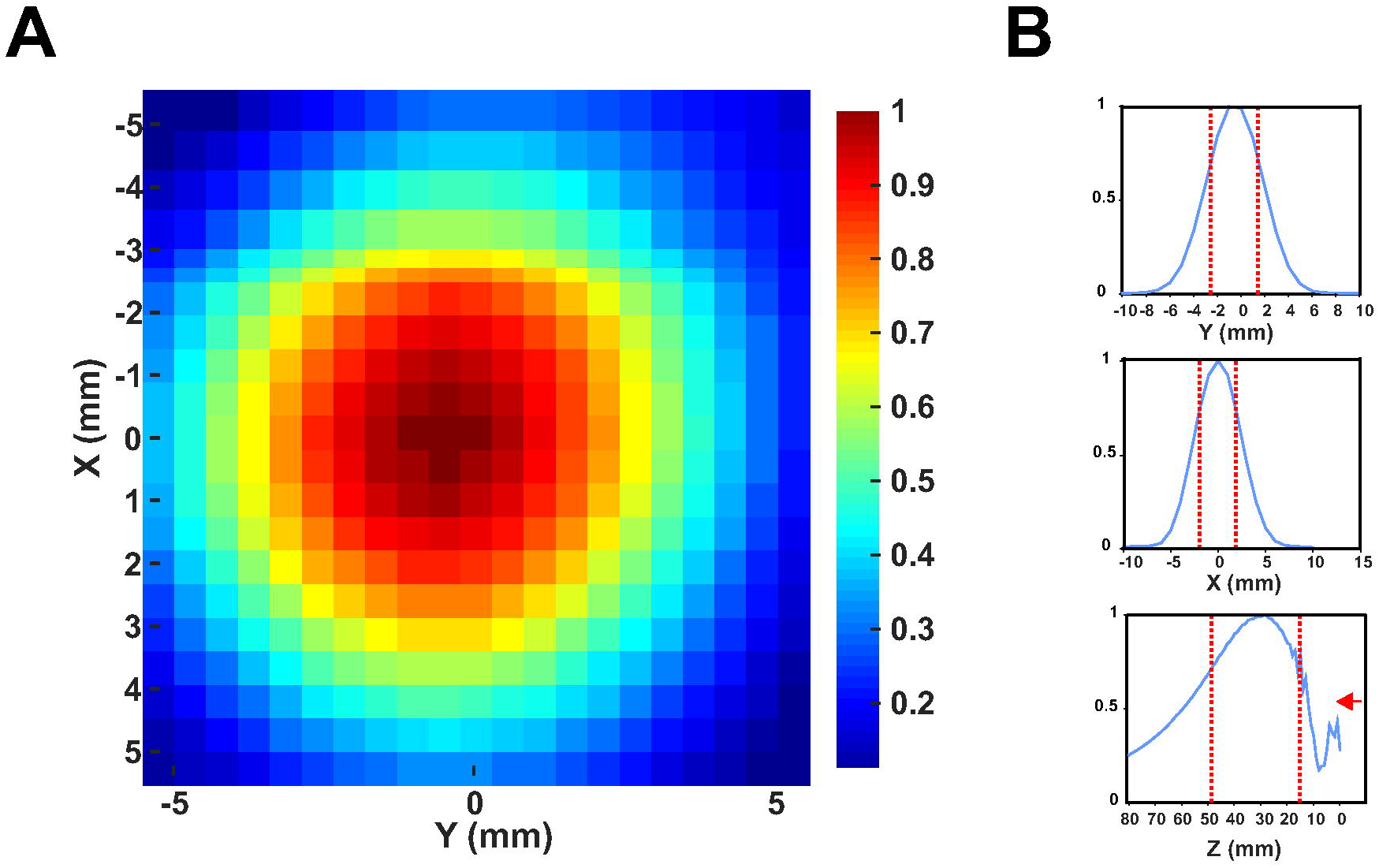
**A**. Pseudocolor XY plot of ultrasound pressure profile normalized to peak pressure. **B.** XYZ line plots of ultrasound pressure profile normalized to peak pressure. Vertical dashed red lines denote -3dB pressure. Note: Red arrow in Z-plot indicates direction of ultrasound from face of transducer (0 mm).

### tFUS targeting

The target for tFUS was chosen based on the isolated thumb fMRI representations found in the first MRI session (Figure 3B). The thumb BOLD representation was loaded into a stereotaxic neuronavigation system (BrainSight; Rogue Research Inc, Montreal, Quebec, CA), and targets were created to guide tFUS based on the strongest BOLD signals in M1 with an approximate depth of ∼ 30 mm (based on the focal length of the transducer) from the scalp on a per subject basis (Figure 3B).

**Figure 3.**
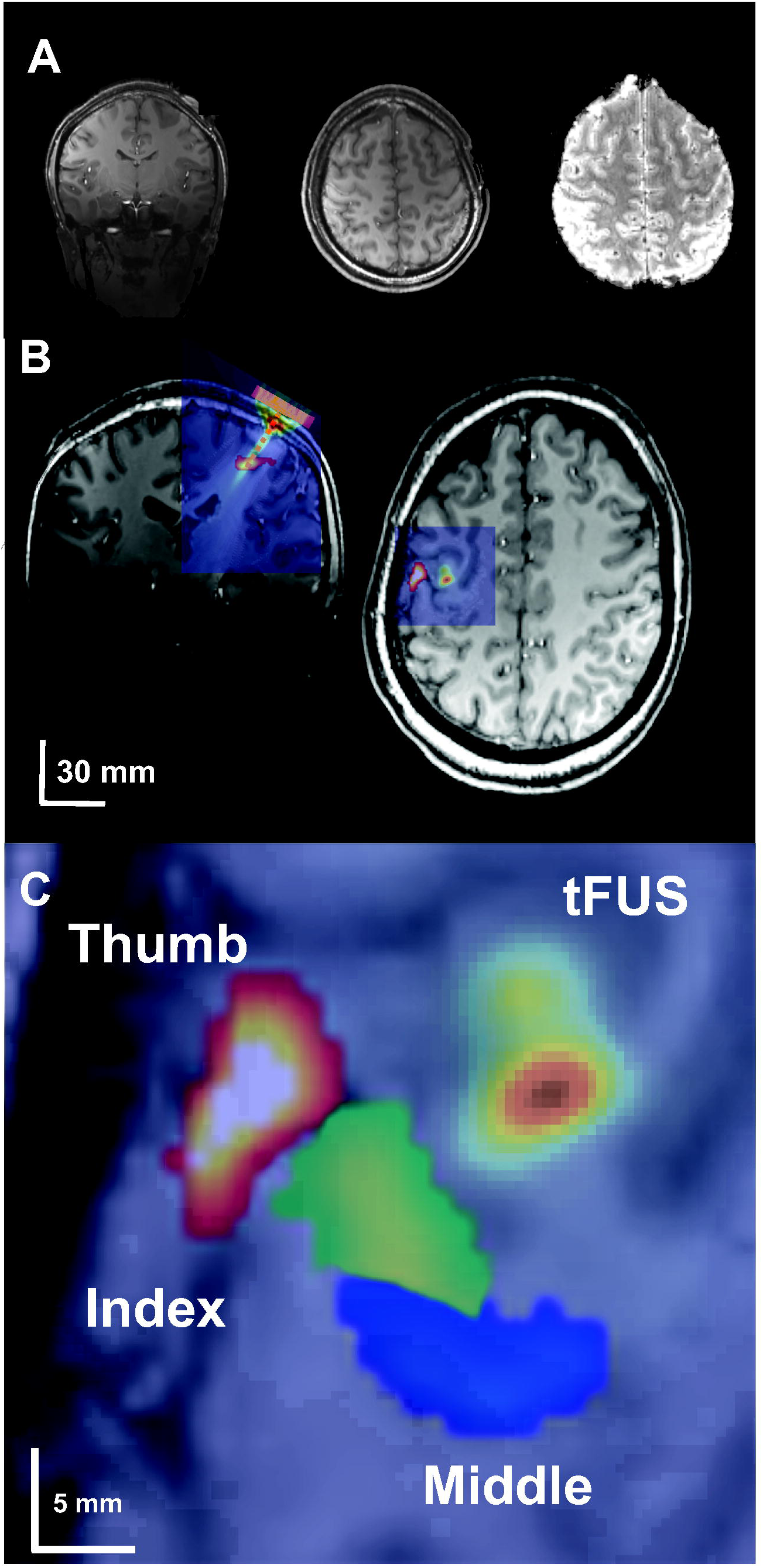
**A**. 7T anatomical T1 (left, middle) and functional EPI image showing ultrasound transducer. **B.** Overlay of functional MRI thumb activation and acoustic model of the ultrasound beam on subject anatomical T1 scan. Note in right image ultrasound beam is purposefully displaced from the fMRI thumb activation to better show relative size compared to fMRI activation. **C.** Blowup of single subject fMRI BOLD finger representations with overlaid acoustic model that is purposefully displaced to show relative size of ultrasound beam to fMRI activations. For experiments, tFUS would have been placed directly over the thumb activation.

### Quantitative modelling of ultrasound wave propagation

To better quantify the intracranial pressure in primary motor cortex from tFUS, a computational model was run to visualize and evaluate the wave propagation of tFUS across an example skull. The model was run using a magnetic resonance (MR) imaging and computerized tomography (CT) dataset taken from the Visible Human Project ® [17]. The transducer was placed on the scalp site overlying the hand knob of the primary motor cortex. Simulations were performed using the k-Wave MATLAB toolbox [18] and modelling parameters and methods are detailed in [14]. The modelled beam is overlaid on a subject an fMRI image of a subject to show the ultrasound beam location relative to the thumb functional activity (Figure 3A) and also to show the lateral resolution of the modelled beam relative to fMRI finger activations (Figure 3C).

### MRI acquisition parameters

All MRI scans were performed at the University of Minnesota’s Center for Magnetic Resonance Research on a 7T Siemens MRI scanner (Siemens Medical Solutions, Erlangen, Germany) using a Nova Medical 1x32 head coil (Wilmington, MA, USA). The fMRI scans were acquired using a gradient echo, echo planar image pulse sequence with the following parameters: TR = 2750ms, TE = 22ms, flip angle = 70, FOV = 192mm x 192mm, number of slices = 108, voxel size = 1.05x1.05x1.05mm^3, iPAT = 3. Additionally, T1 anatomical scans were performed with the following parameters: TR = 3000ms, TE = 3.28ms, flip angle = 6, FOV = 192mm x 216mm, number of slices = 256, voxel size = 1x1x1mm^3.

### BOLD fMRI data analysis

The fMRI data was processed in Analysis of Functional NeuroImages (AFNI) [19]. The data had 3D motion correction, linear and quadratic trends removed, a Gaussian filter with full width half maximum of 3mm applied, slice timing correction, and distortion correction applied. A general linear model analysis was utilized to generate a statistical parametric map with a reference function generated by convolving the hemodynamic response function with the task function. This process was performed for all subjects’ fMRI data to isolate the individual representations of the thumb, index, and middle fingers using a threshold of t = 5 (p = 1e-6 uncorrected). To measure volume changes, a region of interest (ROI) was drawn around the pre-central gyrus (M1) to the depth of the central sulcus. Activated voxels (t = 5; p = 1e-6) in this ROI were used to calculate the activation volume in M1 due to the finger movement being performed for both the tFUS and sham condition. To test for differences between tFUS and sham neuromodulation, the total number of voxels that met this threshold within this ROI was subjected to a paired student’s t test.

For percent signal change analysis, we concentrated on a brain volume at the measured focal volume of the ultrasound beam (see Figure 3). These coordinates were found for each subject and an ROI of 125 mm^3^ (5x5x5 mm) was drawn to encompass partial volume of the ultrasound pressure field. Based upon free water field ultrasound beam measurements, the FWHM volume of the beam was ∼ 230 mm^3^. Percent signal change between tFUS and sham conditions were compared with a paired t test (N = 5). To further investigate the spatial selectivity of the tFUS effect, a 5x5x5 mm ROI was also placed at the region of strongest M1 activations for the index and middle finger representations in each participant to examine if tFUS has effects on these representations despite not being directly targeted for stimulation. Similar group (N=5) paired t-tests were performed separately for the index and middle finger representations.

To test for potential downstream motor network effects as has previously been shown [8], we also examined the effect of tFUS to M1 on the SMA and ipsilateral PMd. The SMA and PMd were defined according to anatomical landmarks. Specifically, SMA included the volume between the precentral and central sulci down to the cingulate sulcus and laterally such that the ROI borders M1 and PMd. The PMd ROI included parts of the superior frontal gyrus and middle frontal gyrus lateral to the SMA and anterior to the pre-central sulcus. Data from the entire scanning session (9 on blocks; thumb, middle and index finger movement; 54 tFUS stimulations) was used in this analysis. We examined both volume and average percent signal from both the SMA and PMd volumes for each participant and each region was tested in a separate group (N=5) paired t-test to assess differences between the tFUS and sham condition.

## Results

### M1 thumb volumes

The application of tFUS at the thumb BOLD representation resulted in larger activation volumes for all five participants (Figure 4A). The group average M1 thumb activation volume was 703±334 mm^3^ for the tFUS condition and 375±167 mm^3^ for the sham condition. The paired t-test revealed a significant increase in BOLD volume for the tFUS condition as compared to sham (t_4_ = 3.01, p = 0.039) (Figure 4B). Table 1 shows the individual subject activation volumes found in M1.

**Figure 4.**
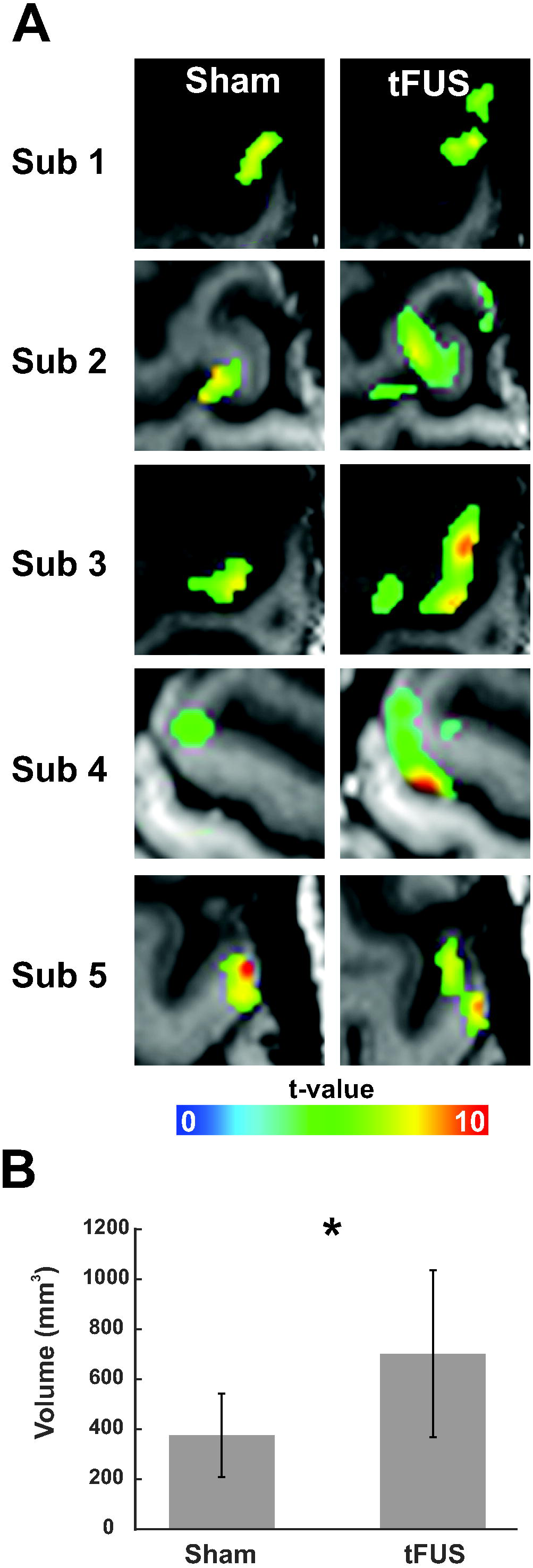
**A**. Individual subject fMRI BOLD thumb activity in primary motor cortex during sham and tFUS neuromodulation. **B.** Group (N = 5) fMRI BOLD M1 volumes for sham and tFUS neuromodulation. * denotes p < 0.05.

The calculated percent changes at the ultrasound beam focus location showed no statistically significant differences between tFUS and sham (Sham: 1.84%±1.36% vs. tFUS: 1.98%±1.17%; t_4_ = 0.7, p = 0.47). See Table 1 for individual participant results.

### Spatial selectivity of tFUS within M1

Based upon previous results that demonstrated high spatial selectivity of ultrasound neuromodulation (Legon et al. 2014) we explored the effect of tFUS on adjacent contiguous volumes within M1. The average Euclidian distance between the center of gravity for the index and middle finger representations were (thumb to index: 10.08 mm ± 5.05 mm; thumb to middle: 10.49 mm ± 6.46 mm). For context, the FWHM lateral resolution of the pressure field is ∼ 5.5 – 6 mm thus the tFUS pressure field can resolve the spatial resolution of the finger representations.

While directing tFUS at the thumb representation we found no differences in activation volumes of the index finger representation (572±999 mm^3^ vs. 665±1428 mm^3^; t_4_ = 0.46, p = 0.67) or the middle finger representation (948±738 mm^3^ vs. 761±793 mm^3^; t_4_ = 0.47, p = 0.80). In addition to BOLD volume changes, we tested for percent signal change and found no differences for either finger representation. The average index finger percent changes were 1.16±1.06% and 2.15±1.79% during the tFUS and sham conditions respectively (t_4_ = 0.46, p = 0.67) and 2.47±1.53% and 2.69±1.95% for the middle finger representation during the tFUS and sham conditions respectively (t_4_ = 0.46, p = 0.67). See Table 1 for individual subject activation volumes and percent changes for the index and middle fingers.

### PMd and SMA

No significant changes were found in SMA between the tFUS and sham conditions for either activation volumes (3191 ± 2966 mm^3^ vs. 2903 ± 2839 mm^3^; t_4_ = 1.35, p = 0.25) or percent signal change (1.92 ± 0.37% vs. 1.87 ± 0.36%; t_4_ = 0.73, p = 0.51). No significant changes were found in PMd between the tFUS and sham condition for activation volumes (202 ± 292 mm^3^ vs. 85 ± 168 mm^3^; t_4_ = 1.86, p = 0.14) or percent signal change (0.65 ± 0.60% vs. 0.66 ± 1.00%; t_4_ = 0.04, p = 0.97).

## Discussion

This is the first study to combine tFUS with 7T fMRI in humans in addition to targeting individual finger representations within M1. The results show that single element 0.5 MHz tFUS targeted at the dominant thumb representation of contralateral M1 increases BOLD activation volumes generated during a cued tapping task. This increase in volume was spatially confined to the sonicated area as it only affected the thumb representation as both adjacent middle and index finger representations did not show any effect. The application of tFUS did not affect percent signal change as compared to sham stimulation and did not have any detectable effect on functionally connected motor regions including the SMA and PMd. These results extend previous results testing the effect of tFUS to elicit a BOLD response [8, 10, 20] and provide for a more detailed perspective on the spatial resolution of tFUS for neuromodulation of individual finger representations within a single gyrus.

The original study by Yoo et al. (2011) in craniotomized rabbits demonstrated 690 kHz focused ultrasound to elicit a BOLD response in M1. The volume of activation was in good spatial approximation with the focus of the pressure field. They did not report any other activation sites suggesting only a local BOLD effect limited to the application site. This BOLD activity was achieved at a relatively low intensity of 3.3 W/cm^2^ and interestingly did not scale with increasing intensity. Double the intensity resulted in a similar increase in percent signal change of around 1.5% from baseline. In Lee et al. (2016) they applied 270 kHz focused tFUS to primary visual cortex (V1) in humans at intensities ranging from ∼ 1 – 10 W/ cm^2^ and reported induced V1 BOLD activity that approximated the pressure field but also reported tFUS to induce activity in functionally connected visual regions. Here, we did not find any evidence for an effect of tFUS on percent signal change in contrast to the above studies or a downstream effect. This is most likely due to differences in experimental design, but also could be related to differences in tFUS parameters. Based upon our previous research that has largely shown inhibition [1, 21], we hypothesized tFUS to also result in inhibition of the BOLD response. As such, we experimentally induced a BOLD signal through a functional motor task and tested the effect of tFUS on this existing signal. It is possible that we did not detect an increase in percent signal change as the motor task had already significantly activated the region and tFUS did not have an additive effect or was undetectable in relation to the strong effect of the motor task. Yoo et al. (2011) reported percent signal changes in the range of 1.5% from ultrasound as compared to resting baseline, though we did not detect any significant increase over our ‘baseline’ that was already at ∼ 1.8 – 2.0 % above rest blocks due to the motor task. It is not clear from Lee et al. (2016) what their signal changes were from their tFUS greater than sham contrast. Unfortunately, we did not test ultrasound during a resting condition in this study to directly compare results with these previous findings for tFUS to induce a BOLD activation. We have previously reported preliminary results in human M1 that showed tFUS to variably induce 3T BOLD activity in 3 of 6 participants though these findings were not robust or statistically significant at the group level [10]. In this study, we were specifically interested in how tFUS affects existing activity and had the specific hypothesis that tFUS would result in inhibition. We assumed that inhibition would translate to a reduction in percent BOLD signal change similar to evoked potential studies where ultrasound attenuated the amplitude of these evoked potentials [1]. However, this was not the case. We found an increase in signal volume and no differences in percent signal change. An increase in signal volume is presumptive of an increase in activity and this could be evidence of the ability of tFUS to produce excitation though it also may be that this increase in volume is a function of increased inhibition. We previously found in Legon et al. (2014) for tFUS to have preferential effects in the gamma band when delivered to primary somatosensory cortex and that this may be a mechanism for the neuromodulatory effect of tFUS. In consideration of the effects found here, a small but very interesting finding in Legon et al. (2014) was for tFUS to increase gamma power when delivered to the precentral gyrus (M1). This somewhat overlooked finding becomes relevant as the gamma frequency band is thought to largely contribute to the BOLD signal [22, 23] and this could explain why we saw an increase in signal volume and would also explain why we did not find an increase in percent signal change. As such, the increase in signal volume we found for all participants in this study could be an indicator of tFUS to preferentially target inhibitory inter-neuronal populations that largely contribute to gamma power [Bartos et al. 2007; Buzsaki and Wang 2012]. This account fits well with data from our lab but is difficult to reconcile with other existing literature that has demonstrated tFUS to motor cortex to elicit peripheral motor responses [20, 24, 25] which would be de facto excitation of pyramidal cells. Here, and in a previous report [10] we do not report any peripheral muscle activity. These discrepancies may be the result of differences in the specific parameters used and/or due to differences in cranial volume or other non-neuronal considerations [26]. In this study, we delivered a total of 54 0.5 second stimulations every 2 TRs (5.5 seconds). This is a higher inter-stimulus interval compared to Yoo et al. (2011) who delivered 3 stimulations every 21 seconds and Lee et al. (2016) that delivered stimulation every 13 seconds though it is unclear how many total sonications were delivered in that study as it is not expressly stated. We employed 500 kHz tFUS which is between what Yoo et al. (2011) and Lee et al. (2016) used though the intensities are similar. These differences may be critical as slight differences in parameters may have a significant impact on the neuronal results as different groups have demonstrated changes in amplitude, duration or duty cycle to affect the neuronal effect [3, 20, 27]. Theoretical accounts of the neuronal effect of ultrasound also predict thresholds for changes in neuronal excitation to inhibition based upon duty cycle and intensity. In the neuronal intramembrane cavitation excitation (NICE) model of the effects of ultrasound our lower duty cycle (36% vs. 50%) may leave us in the transition zone between excitation and inhibition or result only in inhibition [28]. Despite this theoretical model, and the work in small animal models, the effect of tFUS parameters on neuronal excitation in humans is not well understood empirically and indeed the basic putative mechanisms of how mechanical energy affects neuronal excitability is still largely theoretical [28-31]. There is evidence for US to affect certain mechanosensitive channels [32, 33] but the proliferation and density of these channels in human CNS is not well understood and the contribution of these channels to pyramidal excitation and neurovascular coupling is also unclear.

Another important difference between animal studies that show motor excitation and our results is cranial volume. We have previously demonstrated that skull size relative to the ultrasound beam size plays an important role in the intracranial propagation of ultrasound such that smaller skulls or cranial volumes lead to greater interaction of the sound field and higher pressures [14] that could increase the ultrasound effect and produce excitation. Higher amplitude or intensity is theoretically related to excitation [28] and empirical work in oocytes [32] and mice [3] has shown excitation to be a function of amplitude. The waveform we used here measured ∼ 17W/cm^2^ in free water and is estimated from empirical observations through hydrated human skull and through detailed acoustic models to attenuate 4 – 6 times depending on specific properties of the skull [1, 14]. Unfortunately, we were not able to collect computed tomography scans of the subjects here to accurately model and estimate intracranial pressures though the above estimates are in a similar range to previous human studies [1, 34, 35].

In addition to assessing the effect of tFUS on existing BOLD activity, we were also interested in the spatial selectivity of this effect. To examine this, we had participants perform a cued finger tapping task with one of three digits (thumb, index, middle) and only delivered tFUS to the thumb representation during each finger movement. This allowed us to explore the effect of tFUS to not only the targeted thumb region but also on the adjacent non-stimulated index and middle finger regions. We did not find similar index and middle finger volume expansions while tFUS was directed at the thumb representation indicating local spatial effects like those found by [20].

We did not find any evidence that application of tFUS to M1 is able to significantly affect downstream functionally connected regions of the motor system. This finding is at odds with Lee et al. (2016) that reported tFUS directed at V1 to also result in activity in functionally connected regions of the human visual system. Again, differences in experimental design and/or stimulation parameters likely contribute to these differences. The task we used indeed activated both the SMA and the ipsilateral PMd and we do see a weak trend for volume changes in PMd but perhaps the local mechanisms that results in volume increases are limited to the immediate spatial vicinity and are not robust enough to affect downstream regions. One possibility is for the ultrasound effect to be too spatially restricted in that we may have “missed” the targets or not activated enough volume for downstream modulation. Indeed, the effect of non-invasive neuromodulation looks to be spatially and functionally specific as Opitz et al. (2016) showed that depending upon transcranial magnetic stimulation (TMS) current direction to the dorsal lateral pre-frontal cortex different functionally connected networks were activated despite similar spatial locations [36]. As such, due to the spatial restriction of tFUS it is possible that we were not in the ideal spot to effect SMA and PMd activity. It is also possible that again, the motor task sufficiently activated these regions and tFUS did not have an appreciable effect above this level of activity.

Finally, an important consideration when pairing tFUS with MRI and BOLD is for the possibility that the detected response is a result of mechanical energy acting directly on the microvasculature and not on neuronal populations to induce neurovascular coupling. This is likely not the case as pressure levels used here are too low to affect the vasculature. Kaye et al. (2011) demonstrated that focused ultrasound delivered up to 620 W/cm^2^ results in tissue displacement on the order of micrometers, and that this displacement was not detectable in an EPI magnitude MRI image [37].

## Conclusion

This study demonstrated that single element focused ultrasound can be paired with high field 7T fMRI to target individual finger representations within primary motor cortex. With continued research, the pairing of ultrasound with MRI can prove to be a valuable combination for high resolution mapping of discrete brain circuits both cortically and sub-cortically.

## Declarations

### Ethics approval and consent to participate

This study was approved by University of Minnesota’s Institutional Review Board and all Participants gave written informed consent to participate. IRB# 1508M77355

### Consent for publication

Not applicable

### Availability of data and materials

The datasets used and/or analyzed during the current study are available from the corresponding author on reasonable request

### Competing Interests

The authors declare that they have no competing interests

### Funding

Funding for this study was provided by the Department of Rehabilitation Medicine and MNDrive at the University of Minnesota

### Authors’ contributions

LA, JKM and WL were responsible for experimental design, collection, analysis and manuscript preparation. PB was responsible for participant recruitment, collection and manuscript preparation.

## Acknowledgements

Not applicable

